# Brain network dynamics codify heterogeneity in seizure propagation

**DOI:** 10.1101/2021.06.12.448205

**Authors:** Nuttida Rungratsameetaweemana, Claudia Lainscsek, Sydney S. Cash, Javier O. Garcia, Terrence J. Sejnowski, Kanika Bansal

## Abstract

Dynamic functional brain connectivity facilitates adaptive cognition and behavior. Abnormal alterations within such connectivity could result in disrupted functions observed across various neurological conditions. As one of the most common neurological disorders, epilepsy is defined by the seemingly random occurrence of spontaneous seizures. A central but unresolved question concerns the mechanisms by which extraordinarily diverse dynamics of seizures emerge. Here, we apply a graph-theoretical approach to assess dynamic reconfigurations in the functional brain connectivity before, during, and after seizures that display heterogeneous propagation patterns despite sharing similar origins. We demonstrate unique reconfigurations in *globally-defined* network properties preceding seizure onset that predict propagation patterns of impending seizures, and in *locally-defined* network properties that differentiate post-onset dynamics. These results characterize quantitative network features underlying the heterogeneity of seizure dynamics and the accompanying clinical manifestations. Decoding these network properties could improve personalized preventative treatment strategies for epilepsy as well as other neurological disorders.

## Introduction

The human brain is a complex system, where a single brain region interacts with many others and collectively, these interactions give rise to a wide variety of functional connectivity patterns serving rich cognitive functions and adaptive behaviors. A foundation for capturing and understanding such rich and diverse patterns of connectivity is through a graph theoretical framework, where the brain is visualized as a graph or network composed of nodes and edges that represent brain regions and their pairwise associations, respectively *(1–3)*. Architectural features and temporal reconfigurations of the brain’s functional connectivity networks derived within this framework have been associated with several aspects of cognitive functions and development *(1, 4, 5)*. Investigating the temporal evolution of such networks has also provided better insight into the emergence of neural properties such as specialization and efficiency of information processing, learning, and aging *(6–11)*. Applications of network approaches has recently gained popularity particularly in clinical neuroscience for the potential to establish biomarkers of disease onset and progression. Specifically, grounded in graph theory, such techniques can identify abnormal alterations in the dynamic functional brain connectivity that are caused by neurological conditions *(6, 12–16)*.

As one of the most common neurological disorders with roughly 50 million cases world-wide, epilepsy is characterized by its emerging spontaneous seizure activity *(17, 18)*. Critically, one-third of the patients do not respond to medications and rely on alternative interventions such as surgical and neuromodulatory approaches which have variable outcomes and subjective success rates *(19–21)*. Tailoring effective treatment strategies requires biomarkers that can reliably predict not only the temporal complexity but also the dynamical properties of an impending seizure *(22, 23)*. However, seizures are remarkably diverse, making the search for such cues extremely challenging. One way in which the variability across seizures has been addressed is through classification on the basis of the onset regions: *focal seizures* originate from a localized region within one hemisphere while *generalized seizures* begin simultaneously from both hemispheres. A variety of computational techniques in the realm of network science and dynamical systems have been employed to better localize the onset regions and thus improve the precision with which focal and generalized seizures can be identified *(24–26)*. However, localizing onset regions does not fully capture the breadth of dynamics and diversity associated with seizure subtypes. Adding to this complexity, once generated, a focal seizure can remain localized within the same hemisphere (i.e., *focal seizures that remain focal)* or propagate to the other hemisphere (i.e., *focal to bilateral tonic-clonic seizures* or *focal seizures with bilateral spread) (27–29)*. Notably, these subtypes of focal seizures can coexist in a single patient (Fig. 1), where the seizures with bilateral spread generally lead to more severe behavioral and cognitive deficits that could require several minutes to hours for patients to recover from. However, the distinct propagation dynamics exhibited by different seizure types are largely ignored by traditional intervention approaches. Critically, the field currently lacks an objective analytical framework that can be utilized to investigate, understand, and predict the heterogeneity associated with propagation dynamics of seizure activity.

**Fig.1.**
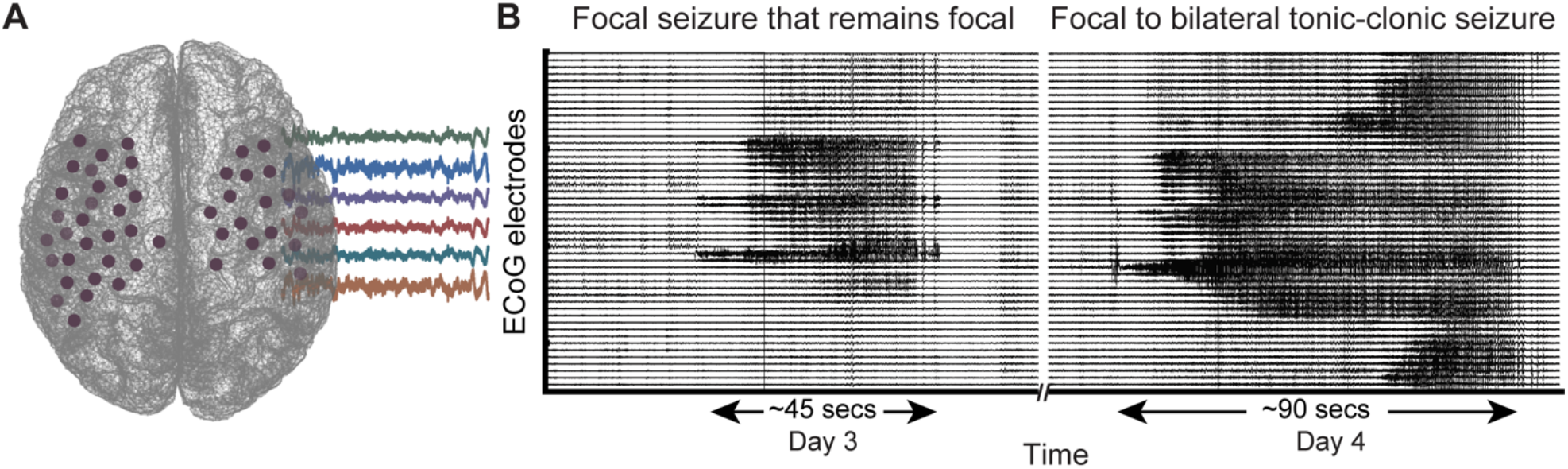
Emergence of distinct seizure propagation patterns in a single patient. **(A)** During a clinical monitoring procedure to identify a seizure onset zone of patients with medication-refractory (drug-resistant) epilepsy, intracranial recording electrodes are implanted. **(B)** Intracranial activity during two sample seizures recorded from a single patient, which exhibit distinct propagation dynamics. On the left, the seizure activity originates from a few electrodes and persists in the localized area within a single hemisphere (i.e., focal seizure that remains focal). On the right, the seizure activity originates from a few electrodes but diffuses bilaterally to involve electrodes in both hemispheres. This type of seizure is known as focal to bilateral tonic-clonic seizure or focal seizure with bilateral spread. Despite their similarly focal origin, these seizure types induce drastically differential clinical manifestations such that focal to bilateral tonic-clonic seizures are associated with more severe cognitive and behavioral deficits. We hypothesize that such heterogeneity in seizure dynamics emerges from distinct and measurable temporal alterations in the functional brain connectivity networks.

Here, we demonstrate that the long-standing challenges associated with the heterogeneity observed across subtypes of epileptic seizures can be addressed through the lens of graph theory. Our study is built upon the idea that the manner in which functional connectivity networks within the brain reconfigure over time carries information concerning the emergent global dynamics and cognitive behaviors that are unique to the underlying neurological processes. Consequently, we probed the time-varying changes within functional networks derived from multiple hours of electrocorticogram (ECoG) recordings across 14 patients as they experienced focal seizures that remain focal or focal to bilateral tonic-clonic seizures. With this analytical framework, we aimed to gain insight into the unique nature of how the heterogenous dynamics associated with different seizure types develop and unfold in the brain. Specifically, we focused on assessing rapid alterations in the architectural attributes that provide quantitative descriptions concerning several aspects of functional connectivity networks of seizures. More specifically, we evaluated these reconfigurations before, during, and after onset of seizures that exhibited drastically different propagation patterns despite sharing similar focal origin.

Our results elucidate key network components that characterize the differential neural dynamics as well as the distinct cognitive and behavioral changes associated with each type of focal seizures. We show that there exist intrinsic network signatures preceding seizure onset that are predictive of the extent to which seizure activity would propagate through the brain. Furthermore, such features emerge several minutes prior to the onset and could, therefore, aid development of successful preventative treatments. Finally, our results reveal differential network characteristics that emerge after seizure onset and characterize the distinct propagation mechanisms of seizure subtypes, suggesting a role of network reconfiguration in regulating termination of seizures. Together, our findings elucidate the association between the evolution of seizures and their underlying network dynamics and offer exciting avenues where graph theoretical measures could be used to guide personalized clinical interventions for neurological disorders such as epilepsy, which displays extensive heterogeneity in its clinical and neurological manifestations across as well as within individual patients.

## Results

### Time-varying functional connectivity networks of focal seizures and interictal activity

To examine the relationship between functional network architecture and propagation mechanisms of focal seizures, we first estimated functional brain connectivity networks from human intracranial recordings of 67 seizures (across 14 patients, 49 focal seizures that remain focal and 18 focal seizures with bilateral spread; Fig. 1B) and 67 interictal periods. For each individual seizure of either type, we epoched a 25-minute segment of electrocorticography (ECoG) data from 15 minutes before to 10 minutes after seizure onset. ECoG data for interictal periods of identical epoch size were chosen with the criterion that such ‘seizure-free’ activity had to take place at least an hour away from an onset and offset of any seizure. Based on these data, we then constructed a series of time-varying connectivity matrices corresponding to each seizure and each interictal period where functional interactions (i.e., connection strength) between the intracranial electrodes were inferred through pairwise cross correlations in sliding 1-second windows *(30–33)*. We then applied a 30-second windowed temporal smoothing procedure such that each of the 25-minute segments of seizure and interictal activity were represented with 98 consecutive functional connectivity matrices (see Materials and Methods and Fig. S1). Based on the graph theoretical framework, these inter-electrode relationships were represented by network *edges* and the electrodes themselves were represented by *nodes* in the corresponding networks of seizures and interictal activity (see Materials and Methods; Fig. 2, A-D).

**Fig.2.**
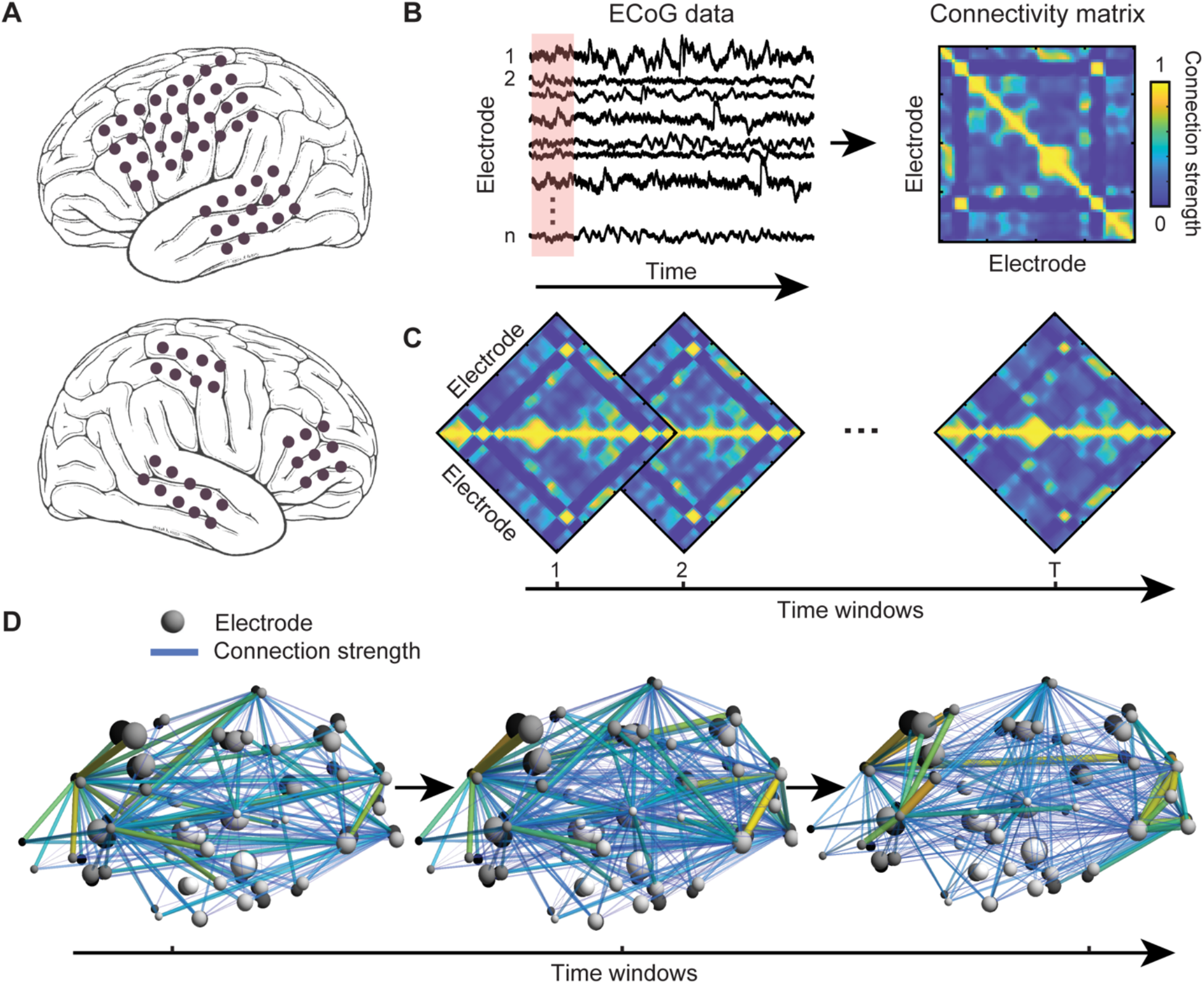
Schematic of graph theoretical analysis of functional brain dynamics. (**A**) Locations of implanted intracranial electrodes of a sample patient. (**B**) We use electrocorticography (ECoG) time-series data from all intracranial electrodes from each patient recorded during a clinical monitoring procedure to locate the seizure onset zone. We estimate the instantaneous functional connectivity of the underlying brain network by computing pairwise correlations of ECoG data across electrodes in a sliding-window manner. The magnitudes of these correlations (restricted between 0 and 1) reflect the strength of connections between each pair of electrodes and are represented by a weighted adjacency or connectivity matrix (see Materials and Methods). (**C**) To investigate time-varying changes in the functional brain connectivity during temporal evolution of each seizure type, we compute a series of connectivity matrices over time and use these as bases to construct functional connectivity networks. (**D**) A schematic of sample constructed networks, consisting of nodes (electrodes) and edges (connection strength). To quantify alterations within these complex networks over time, we evaluate changes of a series of graph theoretical attributes which describe globally- and locally-defined properties of the constructed networks.

To probe the alterations in these functional connectivity networks over time, we extracted a variety of graph theoretical measures which are described in detail in the Materials and Methods section and will be further discussed in the following paragraphs. We assessed and compared how these network attributes evolved during the 25-minute windows corresponding to (i) the dynamics of focal seizures that remained focal (constrained propagation), (ii) the dynamics of focal seizures that became bilateral tonic-clonic seizures (unconstrained propagation), and (iii) the dynamics of ‘seizure-free’ interictal activity. Such analysis approach allowed us to directly test whether there existed unique reconfigurations of network architecture (before, during, and after seizure propagation) that gave rise to the divergent propagation patterns and diverse clinical manifestations exhibited by different subtypes of focal seizures.

### More prominent small-world connectivity links to bilateral spread of seizure activity

Small-world connectivity, characterized by a combination of dense local clustering of connections between neighboring nodes and a short path length between node pairs *(1, 2, 6, 34)*, has been an attractive model for the functional organization of a healthy brain network (see Materials and Methods for more details). In the context of epilepsy, however, increased small-world connectivity has been put forth as a potential driver for the propagation of pathological synchronous activity across brain regions *(32, 35, 36)*. Consequently, we first assessed the clustering coefficients and the characteristic path lengths to examine whether such properties relate to the constrained and unconstrained propagation mechanisms associated with focal seizures that remain focal and focal seizures with bilateral spread, respectively. To accomplish this, we calculated the clustering coefficient and the characteristic path length of each adjacency matrix (i.e., 98 matrices per each of the 25-minute segments of seizure activity). To evaluate these results in light of past studies, we computed averages of these values in a series of consecutive 5-minute windows, separately for each seizure type. This resulted in 3 preictal, 1 ictal (during seizure), and 1 postictal windows (Fig. 3, A and B). A similar analysis was applied to interictal data to estimate baseline values to which the seizure-related network measures could be compared *(37)*.

**Fig 3.**
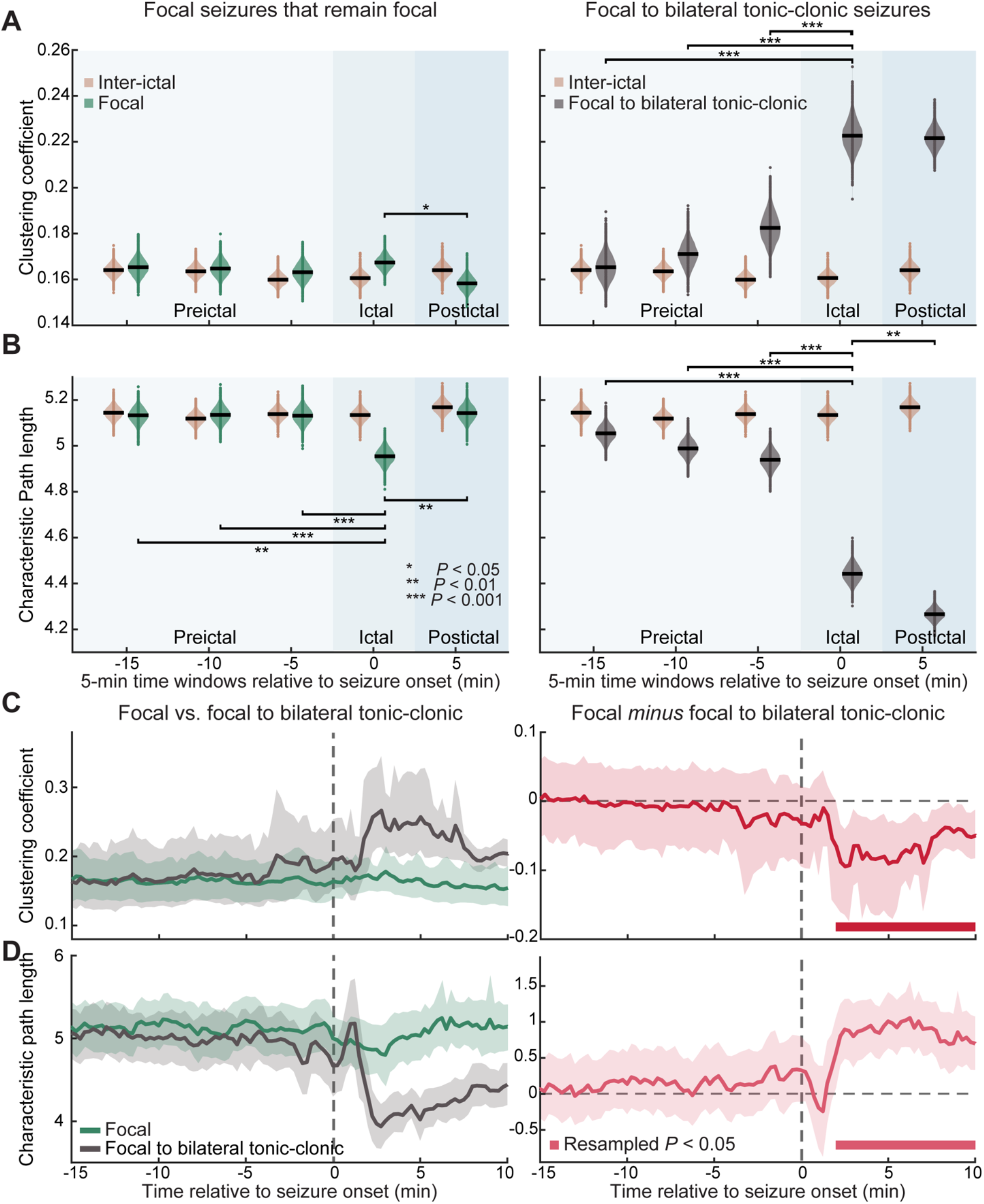
Small-world architecture tracks diffusivity of seizure activity. Focal to bilateral tonic-clonic seizures *(n* = 18) display more prominent small-world connectivity (simultaneous increase in the clustering coefficient and decrease in the characteristic path length) than focal seizures that remain localized within one hemisphere *(n* = 49). (**A**) Averages of the clustering coefficients associated with each seizure type are plotted separately for preictal, ictal (during seizure), and postictal periods. The clustering coefficient of interictal (seizure-free) networks are also plotted as a baseline. (**B**) The characteristic path length is plotted in the same manner. (**C**) The clustering coefficient of focal to bilateral tonic-clonic seizures is higher than that of focal seizures that remain focal, 2-10 minutes after seizure onset. (**D**) The characteristic path length of focal to bilateral tonic-clonic seizures is lower than that of focal seizures that remain focal, 2-10 minutes after seizure onset. Error bars indicate 95% CIs computed by resampling the data distributions. Solid bars show resampled *P* < 0.05.

Our results reveal that both focal seizures that remain focal and focal to bilateral tonic-clonic seizures displayed higher small-world connectivity during ictal periods when compared to seizure-free (interictal) activity as demonstrated by higher clustering coefficients and shorter characteristic path lengths (Fig. 3, A and B). Further analyses demonstrated that the ictal activity associated with focal seizures that remain focal exhibited (i) higher clustering coefficients as compared to postictal period *(P* = 0.03; Fig. 3A, left panel) and (ii) lower characteristic path lengths as compared to both preictal and postictal periods (preictal: *P* = 0.0004, < 0.0001, 0.0002; postictal: *P* = 0.0006; Fig. 3B, left panel). Additionally, we observed similar changes for focal seizures with bilateral propagation where the ictal activity displayed (i) higher clustering coefficients as compared to all the preictal periods (all *P* < 0.0001; Fig 3A, right panel) and (ii) shorter characteristic path lengths as compared to all the preictal periods (all *P* < 0.0001; Fig 3B, right panel). However, unlike the characteristic path lengths associated with the postictal periods of focal seizures with constrained dynamics which returned to the preictal levels, the postictal path length of focal seizures with bilateral spread exhibited a continued decrease *(P* = 0.0006, Fig 3B, right panel). These results supported the more unconstrained diffusivity associated with focal to bilateral tonic-clonic seizures. Notably, the observed differences regarding the manner in which the small-world architecture increased in the networks of focal seizures with constrained and unconstrained dynamics were our first evidence in support of the notion that there may exist network-level signatures that contained information about the distinct propagation mechanisms of focal seizures.

Further, we directly compared the temporal profiles of the small-world architecture for focal seizures that remain localized and focal seizures with bilateral spread. To account for the unequal number of seizure samples of each seizure type, we implemented a bootstrapping procedure and established 95% confidence intervals based on which significant difference was assessed (see Materials and Methods). As expected from Figures 3A-B, we observed that the dynamics of small-world properties differed in a *seizure-type specific manner* only after the onset of seizures. Specifically, the clustering coefficient of focal seizures with bilateral spread was higher than that of focal seizures that remain focal, from 2 to 10 minutes after seizure onset (resampled *P* < 0.05; Fig. 3C). Such differences were accompanied by the shorter characteristic path length associated with focal seizures with bilateral spread (resampled *P* < 0.05 for 2-10 minutes after seizure onset; Fig. 3D). Notably, these observed differences emerged only after the onset and extended well beyond termination of seizures *(36)*, suggesting that focal to bilateral tonic-clonic seizures differentially induced network reorganization that persisted even after the seizure activity ended.

Additionally, after seizure onset, persistent differences in the clustering coefficient and the characteristic path length were also observed between focal seizures with bilateral spread and interictal activity such that focal seizures with bilateral spread displayed more prominent small-world configuration (Fig. S2, right panels). These persistent differences between post-onset activity and interictal periods were, however, not observed in the case of focal seizures that remain focal (Fig. S2, left panels). Together, these findings suggested that the unconstrained propagation dynamics of focal to bilateral tonic-clonic seizures related to an increase in the efficiency of network communication, as illustrated by the increased small-world characteristic shortly after seizure onset. Critically, these observed seizure-type dependent network configurations emerged only after the onset, raising a question whether there also existed unique network alterations at other time points that may contribute to the distinct propagation mechanisms and clinical manifestations associated with each seizure type.

### Alterations in the *local* network connectivity features after the onset reflect heterogeneous dynamics of focal seizures

Given the post-onset differences in the clustering coefficient and the characteristic path length between focal seizures of different propagation mechanisms, we hypothesized that seizure-type dependent network changes should also be observed in other measures of node connectivity patterns such as the degree per node. The degree of a given node is the total sum of edge weights connected to that node. The degree of each individual node therefore reflects the centrality of that node in the network, and the averaged degree per node describes the density of the network. A network with high degree per node is well positioned to optimize integration of information and increase the efficiency of network communication *(34, 37)*. We expected, therefore, that the network nodes after the onset of focal seizures with bilateral spread would be of higher degree on average as compared to those after the onset of focal seizures that remain focal. Supporting our hypothesis, the degree per node associated with focal to bilateral tonic-clonic seizures was found to be higher than that of focal seizures that remain focal for 1.75-10 minutes after the onset (resampled *P* < 0.05, Fig. 4A). Notably, the timing of the sustained differences in the degree per node mirrored that of the clustering coefficient and the characteristic path length, which also extended several minutes beyond seizure termination as each seizure typically lasted between 30 seconds and 3 minutes *(38)*.

**Fig. 4.**
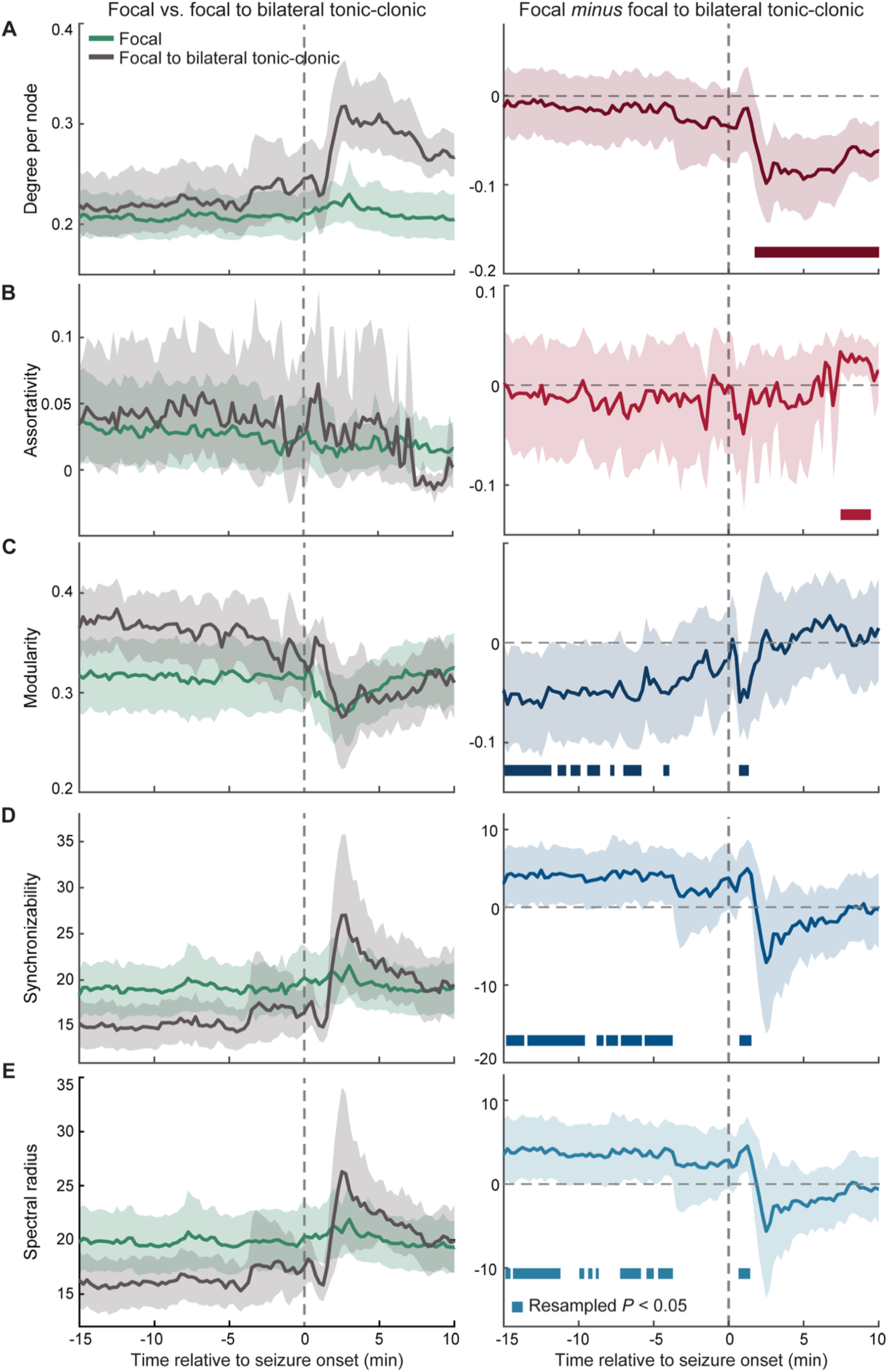
Various features of functional connectivity display distinct temporal changes as a function of seizure propagation dynamics. Left panels illustrate a series of graph theoretical measures computed from networks of focal seizures that remain localized *(n* = 49) and from networks of focal to bilateral tonic-clonic seizures *(n* = 18). The time-varying differences observed in each of these features as a function of seizure types are plotted in the corresponding right panels. (**A**) The degree per node of focal to bilateral tonic-clonic seizures is higher than that of focal seizures that remain focal, 1.75-10 minutes after seizure onset. (**B**) The assortativity, a measure of network robustness, is lower for focal to bilateral tonic-clonic seizures relative to focal seizures that remain focal, 7.5-9.5 minutes after seizure onset. (**C**) The modularity, which captures efficient network integration and global segregation, is higher for focal to bilateral tonic-clonic seizures when compared to focal seizures that remain focal during temporal windows between 14.75 to 3.75 minutes before seizure onset and 0.75-1.50 minutes after the onset. (**D**) The synchronizability, which estimates the propensity of information to diffuse in a network, is higher for focal to bilateral tonic-clonic seizures relative to focal seizures that remain focal during temporal windows between 14.75 to 3.50 minutes before seizure onset and 0.75-1.75 minutes after seizure onset. (**E**) The spectral radius, which relates to the global spread of synchronization in a network, is also higher for focal to bilateral tonic-clonic seizures as compared to focal seizures that remain focal during temporal windows between14.75 to 3.50 minutes before seizure onset and 0.75-1.50 minutes after the onset. Error bars indicated 95% CIs computed by resampling the data distribution. Solid bars show resampled *P* < 0.05.

To further investigate network alterations unique to particular propagation mechanisms of focal seizures, we assessed the assortativity coefficient which measures the propensity of network nodes to connect to other nodes of similar degree *(39, 40)*. In general, network hubs or high-degree nodes of a high-assortativity network are likely to form a highly connected core, surrounded by peripheral nodes with low connectivity. Such configuration renders the network robust against a removal or failure of a single high-degree node. Notably, this core-periphery architecture has been repeatedly observed in functional connectivity networks within the brain *(41, 42)*. In the context of seizures, our results demonstrated that the assortativity coefficient associated with focal to bilateral tonic-clonic seizures was lower than that of focal seizures that remain localized for 7.50-9.50 minutes after seizure onset (resampled *P* < 0.05, Fig. 4B). Additionally, similar patterns of results were observed between focal to bilateral tonic-clonic seizures and interictal activity such that the seizure networks displayed higher degree per node (resampled *P* < 0.05 for 1.75-10 minutes after seizures onset; Fig. S3A) and lower assortativity (resampled *P* < 0.05 for 6.50-7 and 7.50-9.75 minutes after seizures onset; Fig. S3A). However, these network properties did not differ between interictal activity and focal seizures that remain localized.

Importantly, the observed seizure-type differences emerged after the onset of seizures and were contributed by the negative assortativity coefficient that was associated with focal seizures with bilateral propagation. These results suggested that close to seizure termination, the networks of focal seizures with unconstrained dynamics underwent reduced robustness, rendering them more susceptible to network disruptions *(40, 43)*. These findings could potentially account for the more extensive behavioral abnormalities and cognitive deficits often observed after patients experience episodes of focal to bilateral tonic-clonic seizures *(28, 44–46)*.

Thus far, we demonstrated that consistent with the differences in the clustering coefficient and the characteristic path length after seizure onset, the degree per node and the assortativity (i.e., the measures directly derived from local or nodal connectivity), also differed as a function of seizure propagation dynamics. These findings provided better understanding regarding the association between heterogenous propagation mechanisms of seizure activity and the local connectivity within the underlying functional networks. Next, we asked if networks of different seizure types underwent distinct reconfigurations *prior to* seizure onset that shaped the global properties of the networks and ultimately determined the type of propagation dynamics an impending seizure would display.

### Alterations in *global* network features preceding the onset predict propagation dynamics of focal seizures

Building upon the findings presented thus far, we next aimed to quantify the distinct network alterations prior to seizure onset which could differentiate the propagation patterns in a predictive manner. To accomplish this, we assessed network attributes related to various aspects of information processing within a networked system, particularly, the brain connectivity network. Specifically, we focused on three network features: (i) modularity, which represents the tendency of a network to form modules that exhibit strong connectivity within themselves (i.e., strong within-module connectivity) but weak connectivity with other modules in the network (i.e., weak inter-module connectivity) *(47–50)*; (ii) synchronizability, which quantifies how information or activity diffuses in a network *(51, 52)*; and (iii) spectral radius, which describes the speed by which information or activity spreads through a network *(53, 54)*. While modularity has recently been utilized in characterizing the efficiency associated with integration and segregation of information across distributed brain areas, the properties of synchronizability and spectral radius remain relatively unexplored in the context of brain networks. A couple of recent studies, however, have suggested the utility of synchronizability and spectral radius in describing dynamics of seizure activity within the brain *(11)* and the extent of excitability of brain networks, respectively *(10)*. Because modularity, synchronizability, and spectral radius have been associated with different neural processes and are highly sensitive to changes in the network connectivity, we hypothesized that these measures would be powerful markers for prediction of seizure dynamics prior to the onset. As described in the Materials and Methods, each of these attributes relate to overall network architecture and their values may differ across networks with similar distribution of node degrees. Consequently, we characterized modularity, synchronizability, and spectral radius as *global* network features and, in the following, investigated how they change over time as a function of seizure propagation dynamics.

Our results revealed that the information concerning the propagation patterns of focal seizures could be decoded from these global network attributes several minutes prior to seizure onset. Specifically, the modularity preceding the onset of focal seizures with bilateral spread was higher than that of focal seizures that remain localized (resampled *P* < 0.05 for 14.75-11.75, 11.25-9.75, 9.25-8.50, 8.25-8.00, 7.75-7.50, 6.75-5.75, and 4.5-3.75 minutes before seizure onset; Fig. 4C). In addition, the synchronizability associated with focal to bilateral tonic-clonic seizures was lower than that of focal seizures that remain focal (resampled *P* < 0.05 for 14.75-13.50, 13.25-12.25, 12.00-9.50, 9.75-9.50, 8.50-8.25, 8.00-7.25, 7.00-5.75, and 5.50-3.50 minutes before seizure onset; Fig. 4D). This pattern of results was also observed in the spectral radius (resampled *P* < 0.05 for 14.75-14.50, 14.25-11.25, 11.00-11.25, 10.00-9.50, 9.25-8.75, 7.75-7.50, 7.00-5.75, 5.50-5.00, and 4.50-3.50 minutes before seizure onset; Fig. 4E). Additionally, these seizure-type dependent differences in the network modularity, synchronizability, and spectral radius reemerged shortly after seizure onset (resampled P < 0.05 for 0.75-1.50 minutes, 0.75-1.75 minutes, and 0.75-1.50 minutes after seizure onset for modularity, synchronizability, and spectral radius, respectively).

Similar patterns of results were also observed between focal seizures with bilateral spread and interictal activity such that preceding the onset, the seizure networks displayed higher modularity (resampled *P* < 0.05 for 14.75-14, 13.75-12, 10.25-9.50, and 7.75-7.50 minutes before seizure onset; Fig. S3C), lower synchronizability (resampled *P* < 0.05 for 15-13.5, 13.25-9.5, 8.75-8.25, 8-7.25, 7-6.50, and 5.25-3.75 minutes before seizure onset; Fig. S3D), and lower spectral radius (resampled *P* < 0.05 for 14.75-14.50, 14-12.25, 11-9.25, 7.75-7.50, 5.25-4.75, and 4.50-3.75 minutes before seizure onset; Fig. S3E). These results were accompanied by post-onset effects where focal to bilateral tonic-clonic seizures exhibited lower modularity (resampled *P* < 0.05 for 2.50-3 minutes after seizure onset; Fig. S3C), higher synchronizability (resampled *P* < 0.05 for 1-1.50 and 2.50-3 minutes after seizure onset; Fig. S3D), and higher spectral radius (resampled *P* <0.05 for 1-1.50, 2.50-3, and 5.25-5.50 minutes after seizure onset; Fig. S3E). However, focal seizures that remain localized only differed from interictal activity in the measure of modularity such that the modularity of the focal seizures was lower shortly after seizure onset (resampled *P* < 0.05 for 0.75-4.50 minutes after seizure onset; Fig. S3C). These results mimicked the trend observed in the modularity analyses of focal to bilateral tonic-clonic seizures relative to the interictal activity.

Together, our findings illustrated intrinsic topological properties of functional seizure networks preceding the onset that contained information concerning the type of propagation dynamics an impending seizure would display. Importantly, such seizure-type dependent signatures emerged several minutes prior to seizure onset, allowing sufficient time for an effective clinical intervention to be implemented. Furthermore, robust differences in such network attributes reemerged shortly after the onset, confirming the distinct architectural properties associated with the constrained and unconstrained propagation mechanisms of focal seizures. These seizure-type dependent signatures observed post-onset can be used to validate the efficiency of a particular treatment approach in preventing evolution of seizures and may help determine the extent of cognitive and behavioral deficits induced by the residue seizure activity in a scenario where the intervention did not completely eliminate the seizures.

### Complementary temporal reconfigurations within the functional connectivity networks sculpt seizure dynamics

Using a set of graph-theoretical features, we identified reconfigurations in the functional connectivity network that characterized the propagation dynamics of different seizure types. Our results revealed that such distinguishing features can be classified into 2 groups based on the distinct and complementary temporal windows at which the differences in these features emerged as a function of seizure types. The first group of network attributes includes the global features, modularity, synchronizability, and spectral radius, which primarily captures differences between focal seizures with constrained and unconstrained dynamics *prior to* seizure onset (Fig. 5). In contrast, the second group of network properties captured differences across seizure types *after* the onset, reflecting the network reconfigurations induced by distinct propagation mechanisms. Such features include the degree per node, assortativity, clustering coefficient, and characteristic path length (Fig. 5). To further highlight the utility of these features, we evaluated both groups of network measures at a single-seizure level in 3 seizures that had similar onset regions and were recorded from an individual patient (Fig.1 and Fig. S4). Importantly, network features associated with the two seizures (seizure 1 and 3) that remained localized within the left hemisphere exhibited similar temporal patterns which differed from that of the focal seizure with bilateral spread (seizure 2). These results further suggested that network measures could potentially be used to characterize distinct neural dynamics across different types of focal seizures, even on a single-seizure basis.

**Fig. 5.**
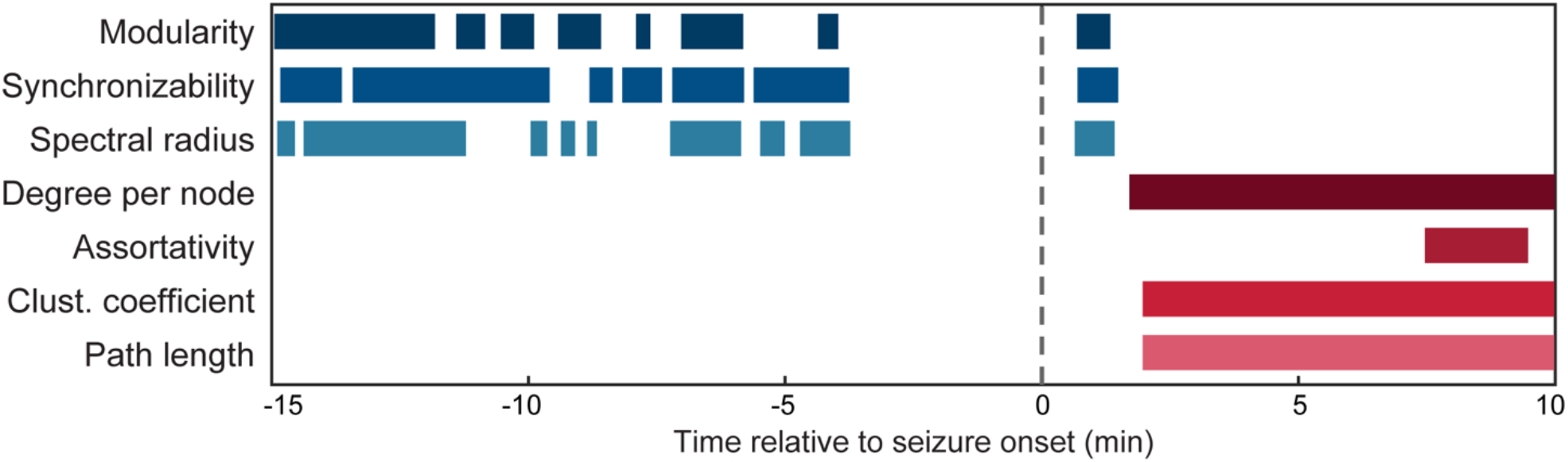
Summary of graph theoretical attributes probed across seizure types. The network features investigated can be categorized into 2 groups based on the temporal windows at which differential changes in these features emerge as a function of seizure propagation patterns. The time windows where such differences are observed are plotted separately for each of the network measures (resampled *P* < 0.05). Global features, i.e., the modularity, synchronizability, and spectral radius, primarily capture network alterations that occur prior to and shortly after seizure onset. In contrast, the degree per node, assortativity, clustering coefficient and characteristic path length characterize post-onset network reconfigurations induced by different types of propagation dynamics.

## Discussion

The present study aimed to investigate if the emergence of heterogeneity in seizure propagation can be understood in terms of network-level changes within the brain before, during, and after the onset. To accomplish this, we evaluated the temporal evolution of a series of graph-theoretical attributes which quantify various aspects of network organization and information processing within complex systems such as the brain. We demonstrated distinct network-level signatures that predicted the extent of diffusion dynamics of an impending seizure as well as isolated architectural changes within the functional connectivity networks that emerged as the seizures terminated. These results advance our understanding of how heterogenous seizure dynamics can arise from similar onset regions. Furthermore, our findings offer exciting avenues where network features may be used to guide clinical diagnosis of seizure subtypes as well as effective intervention strategies to constrain the spread of seizures, thereby minimizing the neurological and cognitive impacts on patients.

### Network alterations track temporal evolution of focal seizures

The brain is an extraordinarily complex system where its microscopic activities constantly fluctuate. However, only the manifestation of such neural processes can be assessed at a macroscopic level through functional neuroimaging techniques. Consequently, features of the functional brain connectivity networks provide a means for characterizing and quantifying the manifestation of the underlying neural activities. Notably, these macroscopic representations as evaluated through the functional connectivity properties, such as the small-world architecture and modular network organization, remain relatively robust despite the fluctuations at the microscopic level. Here, we argue that sustained alterations in the underlying microscopic processes driven by environmental, cognitive, or pathological factors, lead to measurable changes in the observed macroscopic manifestations and consequently the functional connectivity features.

Traditionally, seizures are thought to result from an imbalance between localized excitatory and inhibitory populations in the brain which induces initiation of spontaneous seizure activity *(22, 55)*. On the other hand, the network theory propose instead that seizures are products of aberrant activity in large-scale brain networks *(3, 14, 22, 56)*. In line with this account, we demonstrate that dynamic reconfigurations within the functional connectivity networks during evolution of focal seizures give rise to the heterogeneity observed across seizures. While past work primarily evaluated network properties of epileptic brain by averaging the signals in large discrete time windows, we assess the continuous temporal evolution of network connectivity in combination with resampling statistical tests. This analytical framework enables us to characterize the temporal dynamics of network alterations that underlie the emerging dynamics of seizure activity in an objective and rigorous manner. By examining *globally defined* network features, we observe distinct macroscopic signatures which could predict the extent of diffusivity of seizure propagation minutes prior to the onset. Additionally, we further demonstrate that the heterogeneous dynamics exhibited across seizure types can be characterized by post-onset changes in relatively simplistic features of the functional connectivity networks such as the degree per node. We argue that our methodological approach provides an objective framework not only for better understanding the neural dynamics underlying evolution of seizures but also for determining whether and when a clinical intervention should be implemented to manage and control a spread of an impending seizure.

### Bilateral propagation of focal seizures reflects imbalance in global integration and segregation in the connectivity brain network

We observed that modularity, synchronizability and spectral radius of focal seizures with bilateral spread underwent distinct changes as compared to focal seizures that remain focal and interictal activity. Notably, such differences emerged several minutes prior to seizure onset. In general, modularity, defined as the tendency of a network to separate into high within-connectivity modules, describes the brain’s ability to efficiently integrate information across task-relevant regions while segregating the information across the remaining regions. Likewise, synchronizability and spectral radius also describe global network properties and are specifically used to quantify the ease by with a network can synchronize its activity or processes. Preceding the onset, we reported increased modularity along with decreased synchronizability and spectral radius in focal to bilateral tonic-clonic seizures. Corresponding to a reduced tendency of synchronization and integration within the brain, these observed patterns of the global network properties preceding seizures with greater network diffusion are seemingly counter-intuitive. In light of classical accounts on the mechanistic underpinnings of seizures, we argue that our findings could reflect the chemical or dynamic imbalance within the underlying networks *(22, 57)*. Specifically, due to the microscopic imbalance in excitation and inhibition or the bistability of localized neural dynamics, a neural state can emerge where the connectivity network possesses significantly low integration or compactness, which in turn results in sustained high modularity along with low synchronizability and spectral radius. To regain a more balanced state, it is likely that mechanisms enhancing the connectivity between segregated networks emerge, leading to an over-compensation which results in more unconstrained seizure dynamics. While this is a mere speculation at this point, future studies investigating large-scale non-invasive neuroimaging of epilepsy patients could seek validation and/or refinement to this hypothesis. Notably, we found that such signatures, i.e., increased modularity as well as decreased synchronizability and spectral radius, re-emerged shortly after seizure onset and then disappeared (Fig. 4). It is, therefore, possible that these features capture the manifestations of the regulatory mechanisms that control and prevent the reemergence of seizures. Further support of this conjecture comes from the fact that the modularity of functional connectivity networks also shows an increase in case of focal seizures that remain focal, shortly after seizure onset, when compared to inter-ictal activity (Fig. S3 C), suggesting its link to seizure termination and control mechanisms.

### Increased small-world connectivity induced by the bilateral spread of seizure dynamics

Given its ability to optimize network communication and serve both distributed and specialized information processing, a small-world configuration has been an attractive model for the anatomical and functional structures of the brain *(35, 58)*. In the context of epilepsy, theoretical and empirical work has proposed that the hypersynchronous activity associated with seizures could result from the functional brain networks adopting a configuration that exhibits increased small-world properties *(6, 32, 35)*. We observed an increase in the clustering coefficient and a decrease in the characteristic path length, indicating increased small-world connectivity, following the onset of focal seizures regardless of their propagation mechanisms. Given that the small-world properties capture a balance between the segregation and integration of information within a network, we further hypothesized that the extent of alterations in these measures over time could link to the behavioral and cognitive effects associated with the occurrence of seizures with highly diffused propagation patterns.

Consistent with the post-onset increase in the small-world connectivity, we showed that the seizure-type dependent changes in the network degree per node and assortativity emerged several minutes after seizure termination where focal seizures with bilateral spread display higher degree per node and lower assortativity relative to focal seizures with constrained dynamics. This reduction in the network resilience of focal to bilateral tonic-clonic seizures, as evident by lower assortativity, suggests that bilateral diffusion of seizure activity induced reconfigurations within the functional brain networks that lead to more severe cognitive and behavioral effects observed after termination of focal seizures with unconstrained dynamics. Our findings are also in line with previous reports of decreased assortativity in patients with Alzheimer’s disease *(6, 42)*, and provide early evidence supporting the utility of the assortativity coefficient along with the small-world measures and degree per node in assessing changes in cognitive statuses *(46)*.

### Clinical implications

The heterogeneity of epilepsy is a key confound to disease understanding and development of effective treatments. Here, we demonstrate graph-theoretical features as novel biomarkers that link differential reconfigurations of the functional connectivity networks to the heterogeneity in the emerging seizure dynamics. Specifically, our investigations of the global network dynamics suggest that interventions aiming to contain the spread of seizure activity may wish to situate the brain in a topological state where the modularity is lowered, while the synchronizability and spectral radius are increased. In addition, we also show that the information regarding the propagation patterns of seizures can be decoded through the seizure-type dependent changes in the network properties several minutes before seizure onset allowing sufficient time for an intervention to be implemented. Together, these findings have important clinical implications as monitoring for these early signatures could increase the likelihood of a successful preventative treatment. Furthermore, our results reveal an important link between local network connectivity measures and differential clinical manifestations that are induced by focal seizures with constrained and unconstrained propagation dynamics. Specifically, we demonstrate that the more extensive cognitive and behavioral effects observed after patients undergo focal to bilateral tonic-clonic seizures is associated with the sustained post-onset reconfigurations in the locally defined connectivity features of the underlying networks. These features could also serve as a means to evaluate the effectiveness of an intervention. Future studies that wish to characterize cognitive and behavioral changes induced by neurological disorders may also benefit from evaluating these network properties in relation to performance of patients on various test battery *(6, 59)*. Such analyses could uncover distinct underlying pathophysiological processes that give rise to diverse cognitive and behavioral impairments across disease subtypes and across individuals, thereby improving understanding of the disease heterogeneity *(60–63)*. Critically, such discoveries could guide development of precise and successful clinical interventions that are tailored to diverse neurological conditions. Finally, our single-seizure analyses suggest that network measures could potentially be used to characterize distinct neural dynamics across different types of seizures, even on a single-seizure basis. Such findings provide foundation for future investigation and development of effective personalized seizure treatment.

### Methodological considerations

Given that the electrode placement was determined on a patient-to-patient basis by a neurologist for the purpose of identification of seizure onset zones, the data extracted from these electrodes inevitably provide an incomplete picture of the brain network due to the resulting partial coverage. In addition, the reported lack of differences between focal seizures that remain focal and interictal activity could be partially due to such spatial sampling of the recorded signals. To address this possibility, future studies may benefit from non-invasive recordings where whole-brain dynamics can be simultaneously evaluated. Further, our analyses treated multiple seizures and interictal activity segments from the same patients as independent, and primarily disregarded individual variability in seizure heterogeneity at the patient-level. This analytical choice was made based on traditional methods (e.g., see *(20)*), and careful statistical comparisons were implemented to identify the seizure-type dependent alteration patterns in the functional connectivity networks of seizures. To further extend our findings and improve the specificity of the interpretations, future studies may wish to incorporate patient-level factor in their analytical frameworks.

## Conclusions

In summary, by using a graph theoretical approach, we determined the extent to which distinct emerging dynamics of seizure networks were accounted for by temporal reconfigurations of the underlying functional connectivity. Specifically, we investigated the time-varying changes in network properties associated with focal seizures with constrained and unconstrained propagation patterns. We observed that the network modularity, synchronizability, and spectral radius preceding seizures onset differed between seizures of different propagation dynamics. In addition, the small world measures, degree per node, and assortativity after seizure onset differed as a function of the propagation patterns post seizure onset such that the seizure type dependent differences in these measures reflect the more severe impairments often observed after termination of seizures with bilateral spread. Collectively, our results illustrated a series of network metrics that can be utilized as quantitative biomarkers to distinguish between focal seizures of distinct dynamics on the basis of their propagation patterns as well as the differential extent of cognitive and behavioral effects accompanying the seizures. These results suggested that the networks of focal seizures with unconstrained dynamics undergo early network alterations triggering processes which facilitate the bilateral diffusion of seizure activity. The propagation-type dependent alterations in these metrics were observed again shortly after the onset, suggesting that these measures could also induce regulatory mechanisms necessary for the termination of seizures. Importantly, such findings of network attributes that are unique to seizures of different dynamics several minutes preceding the onset has important clinical implications as tracking fluctuations of these metrics in a time-resolved manner could inform clinicians if an impending focal seizure will diffuse bilaterally. Together, our findings provide objective means to gain better insight into the mechanisms by which seizure dynamics are regulated within the brain and provide exciting avenues where graph theoretical measures could be used to guide personalized clinical interventions.

## Materials and Methods

### Patient information and data acquisition

The seizures analyzed in this study were recorded from 14 patients with medication-refractory epilepsy (Table 1) who underwent a clinical monitoring procedure to locate their seizure onset zone. Clinical electrode implantation, positioning, duration of recordings, and medication schedules were based solely on clinical need as determined by an independent team of clinicians. As indicated in Table 1, seizures analyzed in this study are of two types: 1) seizures that originate from and remain within localized (focal) regions in one hemisphere of the brain (i.e., focal seizures that remain focal); and 2) seizures that originate from focal onset regions in one hemisphere and diffuse bilaterally during the propagation period (i.e., focal to bilateral tonic-clonic seizures or focal seizures with bilateral spread). Patients were implanted with intracranial subdural grids, strips, and depth electrodes for several days in a specialized hospital setting and continuous multichannel electrocorticography (ECoG) data were recorded at a sampling rate of 500 Hz.

**Table 1.** Patient Profiles. Clinical characteristics of the patients. For each patient, we report sex, age at first reported seizures onset, as well as age at the monitoring phase and surgery. We also report the seizure etiology, which was clinically determined through medical history, imaging, and long-term invasive monitoring. Additionally, we indicate the number of observed seizures associated with the two different types of seizures which originated from one hemisphere: focal seizures that remained localized within the same hemisphere (focal seizure; focal) and focal seizures that propagate bilaterally to both hemispheres (focal to bilateral tonic-clonic seizure; focal to bilateral). Surgical outcome (outcome) was based on Engel score: seizure freedom to no improvement (I-V), and no follow-up (NF). Legend: M = male; F = female; MTS = Mesial Temporal Sclerosis; n.a = not applicable.

| Patient | Sex | Age at onset/<br>surgery | Etiology | Seizure Type (#) | Seizure Onset<br>Zone | Resection Areas | Outcome |
| --- | --- | --- | --- | --- | --- | --- | --- |
| 1 | F | 15/46 | Dysplasia | focal to bilateral (5) | Anterior<br>temporal | Right<br>anterior temporal | I |
| 2 | F | 42/55 | n.a | focal to bilateral (3) | Temporal | None | I |
| 3 | F | 17/45 | n.a | focal (1);<br>focal to bilateral (2) | Temporal | None | n.a |
| 4 | M | 8/23 | n.a | focal (10) | Frontal | None | III |
| 5 | M | 14/35 | n.a | focal (9);<br>focal to bilateral (2) | Temporal | Right<br>anterior temporal | n.a |
| 6 | F | 12/32 | n.a | focal (15) | Temporal | Right<br>anterior temporal | II |
| 7 | F | 7/23 | n.a | focal (6) | Frontal | Left frontal | IV |
| 8 | F | 10/27 | n.a | focal (1) | Unknown | Left frontal | IV |
| 9 | F | 8/19 | MTS | focal (1) | Anterior<br>temporal | Left<br>anterior temporal | III |
| 10 | F | 14/31 | n.a | focal to bilateral (2) | Temporal | Right<br>anterior temporal | I |
| 11 | F | 1/21 | Stroke | focal (2) | Temporal | Left temporal | IV |
| 12 | F | 9/42 | n.a | focal (2) | Frontal | None | II |
| 13 | M | 39/47 | n.a | focal to bilateral (3) | Posterior<br>temporal | Right temporal | I |
| 14 | F | 50/59 | n.a | focal (2);<br>focal to bilateral (1) | Posterior<br>temporal | Left temporal | I |

Only seizures with an obvious ictal onset were selected for analysis. Experienced epileptologists, blind to this study, identified the seizure onset regions, seizure types, and onset time through inspection of the ECoG recordings, referral to the clinical report, and clinical manifestations recorded on video. A total of 67 seizures (49 focal seizures that remain focal and 18 focal to bilateral tonic-clonic seizures) were analyzed. We note that, multiple seizures from the same patients were treated as independent (see similar methods in *(20)*). All patients were enrolled after informed consent was obtained and approval was granted by local Institutional Review Boards (IRB) at Massachusetts General Hospital (MGH) according to National Institutes of Health (NIH) guidelines.

### Data preprocessing

For each of these seizures, we considered ECoG data of the duration of 15 minutes before and 10 minutes after the seizure onset. Each of these 25-min data segments only contained one seizure. For comparison with relatively ‘seizure-free’ activity, we extracted an equal number of interictal activity windows with the same duration. Interictal windows were selected from ECoG recordings at least an hour away from an onset and offset of any seizure. The data were band-pass filtered between 1 to 70 Hz, and notch filtered at 60 Hz to exclude potential powerline interference. A common reference was used for data analysis and the reference electrode in each case was located far from the area of recording making the introduction of spurious correlation or elimination of actual correlation between cortical regions unlikely *(64)*.

### Functional connectivity networks

To evaluate functional connectivity representations associated with the temporal evolution of seizures, we employed complex network analysis. Originated from the mathematical study of networks known as graph theory, such analytical framework represents a real-world complex system as a network that is composed of a collection of nodes with edges connecting different node pairs. Here, we constructed functional brain connectivity networks where edges represented cross correlations between pairs of recording electrodes over time.

Specifically, we computed symmetric functional connectivity *C*_*ij*_ between two regions of the brain *i* and *j* as an averaged correlation of the neural signals recorded by the intracranial electrode contacts of those regions. To extract (at least approximately) the stationary aspects of ECoG data, we divided each of the 25-min ECoG data segments into consecutive 1-s windows, where each window overlapped the previous window by 0.5 seconds *(56, 65)*. The correlation was calculated within each of these 1-s segments. To account for noise, we applied a temporal smoothing to these correlation values by averaging consecutive 30 seconds windows such that a total of 98 correlation values representing the functional connectivity of 98 temporal windows were generated from each 25-min ECoG data segment. Note that different temporal smoothing parameters can be use without affecting the overall patterns of results, although a value too large may reduce the temporal precision of the observations (Fig. S1).

All correlation values were bounded between −1 and +1. Negative correlation values implying long range inhibitions were then set to zero, as within our modelling framework and in line with previous studies *(19, 66)*, we do not consider the contribution of long range direct inhibitory connections to the simulation of the epileptogenic effect. Temporal evolution of these correlations or adjacency matrices reflects the time-varying connectivity dynamics of the functional brain network from which the ECoG data were recorded.

### Graph theoretical network analysis

For each seizure, we constructed a series of weighted, symmetric (undirected) adjacency matrices (connectivity matrices) *C* representing functional connectivity networks across all recording electrodes. From these networks, we computed a series of graph theorical network measures (described below) as a function of seizure types to quantify changes in network dynamics associated with evolution of focal seizures with constrained (focal seizure that remain focal) and unconstrained propagation mechanisms (focal to bilateral tonic-clonic seizures). We used the various Brain Connectivity Toolbox functions implemented in MATLAB (R2020; MathWorks) for our computation of these network features unless noted otherwise.

#### Assessing small-world architecture

In general, a network can range from completely regular where each node connects to its nearest neighbors to fully random where node pairs are connected randomly with some probability *(52, 67–69)*. Within this spectrum lies a small-world architecture which is characterized by a combination of dense local clustering of connections between neighboring nodes (like regular networks) and a short path length between distant node pairs due to the existence of relatively few long-range connections (like random networks) *(1, 2, 6, 34)*. This architectural scheme is known to facilitate both specialized and distributed information processing in a cost-effective manner, and thus has been an attractive model for the functional organization of a healthy brain network. Mathematically, small-world architecture is characterized by high clustering coefficient and low Characteristic path length as compared to a random network.

#### Clustering coefficient

Clustering coefficient is a measure of local connectedness of a network and has been used to describe the segregation of information in brain networks. The clustering coefficient is calculated as the ratio between the number of triangles present around a node and the maximum number of triangles that could possibly be formed around that node *(16, 70)*. For a given node X and any other two nodes Y and Z within the network, a triangle around X represents a scenario where X, Y, and Z all have a connectivity value of 1 with one another. We used the Brain Connectivity Toolbox function *clustering_coef_wu* for the calculation of clustering coefficient.

#### Characteristic path length

Characteristic path length describes the averaged minimum distance between all pairs of nodes in a network and has been shown to associate with the integration of information within the brain network. The minimum path length between a pair of network nodes represents the shortest route between them through a combination of network edges. We calculated characteristic path length using the Brain Connectivity Toolbox function *charpath (37)*.

In the context of epilepsy research, it has been suggested that increased small-world architecture, i.e., increase in clustering coefficient and decrease in characteristic path length, could facilitate synchronization of seizure activity from the onset zones to other parts of the brain *(32, 35, 36)* and therefore, we considered small-word configuration as an important network attribute to differentiate the dynamics of different seizure subtypes.

#### Degree per node

Degree of a node represents the total sum of edge weights connected to a node in the network. To compare seizures across individuals who had different number of implanted electrodes, we computed the average degree per node which represents on an average, the total sum of edge weights connected to a node in the network. A high average degree per node indicates a large number of connections and this measure represents the ‘wiring cost’ of the network. A network with high degree per node is well positioned to optimize integration of information and increase the efficiency of network communication (31, 34).

For a given node *i*, the degree is defined as 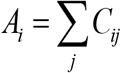 where, C represents the connectivity matrix.

Then, degree per node is calculated as 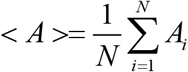 where, N represents the total number of nodes in the network.

#### Assortativity

Assortativity measures the propensity of nodes to connect to others with similar degree and is calculated as a correlation coefficient between the degrees of all the nodes *(39)*. A positive assortativity value indicates that nodes tend to link to other nodes with similar degree, whereas a negative value indicates connected nodes with dissimilar degree. Networks with high assortativity tend to make a highly connected core of network hubs. Functional brain networks have been shown to display such architecture with highly connected hub regions or core surrounded by low-connectivity peripheral nodes. Assortativity quantifies network robustness as a removal or failure of a single high-degree node would induce greater impact on communication efficiency of a network with low assortativity than on a network with high assortativity. By examining the measure of assortativity across seizure subtypes, we could evaluate whether there existed a relationship between different propagation mechanisms and the extent of network resilience. We calculated assortativity using the Brain Connectivity Toolbox function *assortativity_wei (37)*.

#### Modularity

Modularity describes the extent to which a graph can be divided into clearly separated communities (i.e., subgraphs or modules). Each module contains several interconnected nodes, and there are relatively few connections between nodes of different modules. In the context of brain networks, modularity has been used to describe and quantify efficient integration and segregation of information across distributed sets of brain regions as a function of cognitive task demands *(5, 71)*. We used the Brain Connectivity Toolbox function *modularity_und* to compute modularity of functional brain networks *(48, 72)*.

#### Synchronizability

Synchronizability relates to the viability of synchronized dynamics within a network. Particularly in the context of epilepsy, relatively larger value of synchronizability has been associated with greater ease for neural populations to synchronize their dynamics *(11)*. Synchronizability (***S***) is calculated as the ratio of the second smallest and the largest eigenvalue of the Laplacian matrix (***L***), which is computed as the difference between the diagonal matrix of node strength (total degree) and the adjacency matrix such that ***L = D – C***. Thus, synchronizability estimates the spread of the eigenvalues of the network Laplacian and is computed as 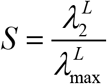 where 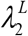 and 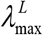 denote the second smallest and the largest eigenvalue of ***L***, respectively.

#### Spectral radius

Spectral radius is a global measure of network structure that is related to the spread of synchronization in a network *(10, 53, 73)*. Computed as the largest eigenvalue of the connectivity matrix (C), spectral radius reflects the critical coupling strength required to synchronize the system *(74)*. As such, spectral radius represents the principal component of the system and contains information about structural characteristics as well as dynamical behavior and stability of the underlying network *(75–77)*. In the network based models of brain dynamics, spectral radius has been associated with the ease with which the system can be transitioned into an excited state *(10)*.

## Statistical analysis

To compare the computed network measures as a function of seizure types in a time-resolved manner and to assess significant differences, we performed a bootstrapping procedure and established 95% confidence intervals for each corresponding measure. For each of the bootstrapping iterations, we performed resampling with replacement at the level of individual seizures and computed averages for comparison of interest (e.g., the clustering coefficient of focal seizures that remain focal vs. the clustering coefficient of focal to tonic-clonic seizures). We performed 10,000 bootstrapping iterations in order to achieve the confidence intervals reported (CIs) for each comparison. Note that this approach constrains the resolution of *P* values to a lower limit of *P* ≤ 0.0001. We generated permuted null distributions of each complex network measure for each individual seizure and each time point. For tests comparing a bootstrapped distribution against 0, *P* values were computed by conducting 2 one-tailed tests against 0 (e.g., mean[difference in clustering coefficients < 0] and mean[difference in clustering coefficients > 0] and doubling the smaller *P* value)

## Acknowledgments

We wish to thank Pariya Salami and Mia Borzello for assistance with data preprocessing. We are also grateful to the clinical team, technicians, and our research participants for their help in making this research possible.

## Author contributions

Conceptualization: NR, KB. Methodology: NR, CL, JOG, KB. Data Curation: SSC. Formal Analysis: NR. Visualization: NR, KB. Supervision: JOG, TJS, KB. Funding Acquisition: SSC, TJS, JOG, KB. Writing – original draft: NR, JOG, KB. Writing – review & editing: CL, SSC, TJS.

## Data and materials availability

All data needed to evaluate the conclusions in the paper are present in the main text and/or the Supplementary Materials. The seizure data analyzed in this study maybe requested from the authors. The data are not publicly available as they contain information that could compromise privacy of the research participants.

## References

1. E. Bullmore, O. Sporns, Complex brain networks: Graph theoretical analysis of structural and functional systems. Nat. Rev. Neurosci. 10, 186–198 (2009).

2. D. S. Bassett, O. Sporns, Network neuroscience. Nat. Neurosci. 20, 353–364 (2017).

3. R. C. Wykes, H. M. Khoo, L. Caciagli, H. Blumenfeld, J. Kapur, J. M. Stern, A. Bernasconi, N. Bernasconi, Wonoep appraisal: Network concept from an imaging perspective. 60, 1293–1305 (2019).

4. J. D. Medaglia, M.-E. Lynall, D. S. Bassett, Cognitive network neuroscience. J. Cogn. Neurosci. 27, 1471–1491 (2015).

5. D. Godwin, R. L. Barry, R. Marois, Breakdown of the brain’s functional network modularity with awareness. Proc. Natl. Acad. Sci. U. S. A. 112, 3799–3804 (2015).

6. C. J. Stam, Modern network science of neurological disorders. Nat. Rev. Neurosci. 15, 683–695 (2014).

7. T. Proix, F. Bartolomei, M. Guye, V. K. Jirsa, Individual brain structure and modelling predict seizure propagation. Brain. 31, 13292–300 (2017).

8. B. G. Y. Ridley, C. Rousseau, J. Wirsich, A. Le Troter, E. Soulier, S. Confort-Gouny, F. Bartolomei, J. P. Ranjeva, S. Achard, M. Guye, Nodal approach reveals differential impact of lateralized focal epilepsies on hub reorganization. Neuroimage. 118, 39–48 (2015).

9. J. O. Garcia, A. Ashourvan, S. Muldoon, J. M. Vettel, D. S. Bassett, Applications of community detection techniques to brain graphs: Algorithmic considerations and implications for neural function. Proc IEEE Inst Electr Electron Eng. 106, 846–867 (2018).

10. K. Bansal, J. D. Medaglia, D. S. Bassett, J. M. Vettel, S. F. Muldoon, Data-driven brain network models differentiate variability across language tasks. PLoS Comput. Biol. 14, 1–25 (2018).

11. A. N. Khambhati, K. A. Davis, T. H. Lucas, B. Litt, D. S. Bassett, Virtual Cortical Resection Reveals Push-Pull Network Control Preceding Seizure Evolution. Neuron. 91, 1170–1182 (2016).

12. D. S. Bassett, E. Bullmore, B. A. Verchinski, V. S. Mattay, D. R. Weinberger, A. Meyer-Lindenberg, Hierarchical organization of human cortical networks in health and Schizophrenia. J. Neurosci. 28, 9239–9248 (2008).

13. B. C. Bernhardt, L. Bonilha, D. W. Gross, Network analysis for a network disorder: The emerging role of graph theory in the study of epilepsy. Epilepsy Behav. 50, 162–170 (2015).

14. W. Stacey, M. Kramer, K. Gunnarsdottir, J. Gonzalez-Martinez, K. Zaghloul, S. Inati, S. Sarma, J. Stiso, A. N. Khambhati, D. S. Bassett, R. J. Smith, V. B. Liu, B. A. Lopour, R. Staba, Emerging roles of network analysis for epilepsy. Epilepsy Res. 159, 106255 (2020).

15. U. Braun, A. Schaefer, R. F. Betzel, H. Tost, A. Meyer-Lindenberg, D. S. Bassett, From maps to multi-dimensional network mechanisms of mental disorders. Neuron. 97, 14–31 (2018).

16. L. C. Huang, P. A. Wu, S. Z. Lin, C. Y. Pang, S. Y. Chen, Graph theory and network topological metrics may be the potential biomarker in Parkinson’s disease. J. Clin. Neurosci. 68, 235–242 (2019).

17. F. Mormann, T. Kreuz, C. Rieke, R. G. Andrzejak, A. Kraskov, P. David, C. E. Elger, K. Lehnertz, On the predictability of epileptic seizures. Clin. Neurophysiol. 116, 569–587 (2005).

18. P. Jiruska, M. de Curtis, J. G. R. Jefferys, C. A. Schevon, S. J. Schiff, K. Schindler, Synchronization and desynchronization in epilepsy: Controversies and hypotheses. J. Physiol. 591, 787–797 (2013).

19. N. Sinha, J. Dauwels, M. Kaiser, S. S. Cash, M. B. Westover, Y. Wang, P. N. Taylor, Predicting neurosurgical outcomes in focal epilepsy patients using computational modelling. Brain. 140, 319–332 (2017).

20. L. E. Martinet, M. A. Kramer, W. Viles, L. N. Perkins, E. Spencer, C. J. Chu, S. S. Cash, E. D. Kolaczyk, Robust dynamic community detection with applications to human brain functional networks. Nat. Commun. 11, 1–13 (2020).

21. P. Salami, N. Peled, J. K. Nadalin, L. E. Martinet, M. A. Kramer, J. W. Lee, S. S. Cash, Seizure onset location shapes dynamics of initiation. Clin. Neurophysiol. 131, 1782–1797 (2020).

22. L. Kuhlmann, K. Lehnertz, M. P. Richardson, B. Schelter, H. Zaveri, Seizure prediction -- ready for a new era. Nat. Rev. Neurol. 14, 618–630 (2018).

23. C. Lainscsek, N. Rungratsameetaweemana, S. S. Cash, T. J. Sejnowski, Cortical chimera states predict epileptic seizures. Chaos. 29 (2019), doi:10.1063/1.5139654.

24. V. K. Jirsa, W. C. Stacey, P. P. Quilichini, A. I. Ivanov, C. Bernard, On the nature of seizure dynamics. Brain. 137, 2210–2230 (2014).

25. M. L. Saggio, D. Crisp, J. M. Scott, P. Karoly, L. Kuhlmann, M. Nakatani, T. Murai, M. Dümpelmann, A. Schulze-Bonhage, A. Ikeda, M. Cook, S. V. Gliske, J. Lin, C. Bernard, V. Jirsa, W. C. Stacey, A taxonomy of seizure dynamotypes. Elife. 9, 1–56 (2020).

26. A. Li, B. Chennuri, S. Subramanian, R. Yaffe, S. Gliske, W. Stacey, R. Norton, A. Jordan, K. A. Zaghloul, S. K. Inati, S. Agrawal, J. Haagensen, J. Hopp, C. Atallah, E. Johnson, N. Crone, W. S. Anderson, Z. Fitzgerald, J. Bulacio, J. T. Gale, S. V Sarma, J. Gonzalez-Martinez, Using network analysis to localize the epileptogenic zone from invasive EEG recordings in intractable focal epilepsy. Netw. Neurosci. 1, 222–241 (2018).

27. R. S. Fisher, J. H. Cross, C. D’Souza, J. A. French, S. R. Haut, N. Higurashi, E. Hirsch, F. E. Jansen, L. Lagae, S. L. Moshé, J. Peltola, E. Roulet Perez, I. E. Scheffer, A. Schulze-Bonhage, E. Somerville, M. Sperling, E. M. Yacubian, S. M. Zuberi, Instruction manual for the ILAE 2017 operational classification of seizure types. Epilepsia. 58, 531–542 (2017).

28. R. S. Fisher, J. H. Cross, J. A. French, N. Higurashi, E. Hirsch, F. E. Jansen, L. Lagae, S. L. Moshé, J. Peltola, E. Roulet Perez, I. E. Scheffer, S. M. Zuberi, Operational classification of seizure types by the International League Against Epilepsy: Position Paper of the ILAE Commission for Classification and Terminology. Epilepsia. 58, 522–530 (2017).

29. H. Blumenfeld, G. I. Varghese, M. J. Purcaro, J. E. Motelow, M. Enev, K. A. McNally, A. R. Levin, L. J. Hirsch, R. Tikofsky, I. G. Zubal, A. L. Paige, S. S. Spencer, Cortical and subcortical networks in human secondarily generalized tonic-clonic seizures. Brain. 132, 999–1012 (2009).

30. S. P. Burns, S. Santaniello, R. B. Yaffe, C. C. Jouny, N. E. Crone, G. K. Bergey, W. S. Anderson, S. V. Sarma, Network dynamics of the brain and influence of the epileptic seizure onset zone. Proc. Natl. Acad. Sci. U. S. A. 111, E5321–E5330 (2014).

31. J. C. Reijneveld, S. C. Ponten, H. W. Berendse, C. J. Stam, The application of graph theoretical analysis to complex networks in the brain. Clin. Neurophysiol. 118, 2317–2331 (2007).

32. M. A. Kramer, S. S. Cash, Epilepsy as a disorder of cortical network organization. Neuroscientist. 18, 360–372 (2012).

33. M. A. Kramer, U. T. Eden, K. Q. Lepage, E. D. Kolaczyk, M. T. Bianchi, S. S. Cash, Emergence of persistent networks in long-term intracranial eeg recordings. J. Neurosci. 31, 15757–15767 (2011).

34. E. Bullmore, O. Sporns, The economy of brain network organization. Nat. Rev. Neurosci. 13, 336–349 (2012).

35. S. C. Ponten, F. Bartolomei, C. J. Stam, Small-world networks and epilepsy: Graph theoretical analysis of intracerebrally recorded mesial temporal lobe seizures. Clin. Neurophysiol. 118, 918–927 (2007).

36. M. A. Kramer, U. T. Eden, E. D. Kolaczyk, R. Zepeda, E. N. Eskandar, S. S. Cash, Coalescence and fragmentation of cortical networks during focal seizures. J. Neurosci. 30, 10076–10085 (2010).

37. M. Rubinov, O. Sporns, Complex network measures of brain connectivity: Uses and interpretations. Neuroimage. 52, 1059–1069 (2010).

38. S. Jenssen, E. J. Gracely, M. R. Sperling, How long do most seizures last? A systematic comparison of seizures recorded in the epilepsy monitoring unit. Epilepsia. 47, 1499–1503 (2006).

39. M. E. J. Newman, Assortative Mixing in Networks. Phys. Rev. Lett. 89, 1–4 (2002).

40. R. Noldus, P. Van Mieghem, Assortativity in complex networks. J. Complex Networks. 3, 507–542 (2014).

41. S. Lim, F. Radicchi, M. P. van den Heuvel, O. Sporns, Discordant attributes of structural and functional brain connectivity in a two-layer multiplex network. Sci. Rep. 9, 1–13 (2019).

42. W. de Haan, Y. A. L. Pijnenburg, R. L. M. Strijers, Y. van der Made, W. M. van der Flier, P. Scheltens, C. J. Stam, Functional neural network analysis in frontotemporal dementia and Alzheimer’s disease using EEG and graph theory. BMC Neurosci. 10, 1–12 (2009).

43. D. Sone, H. Matsuda, M. Ota, N. Maikusa, Y. Kimura, K. Sumida, K. Yokoyama, E. Imabayashi, M. Watanabe, Y. Watanabe, M. Okazaki, N. Sato, Graph theoretical analysis of structural neuroimaging in temporal lobe epilepsy with and without psychosis. PLoS One. 11 (2016), doi:10.1371/journal.pone.0158728.

44. A. T. Berg, S. F. Berkovic, M. J. Brodie, J. Buchhalter, J. H. Cross, W. Van Emde Boas, J. Engel, J. French, T. A. Glauser, G. W. Mathern, S. L. Moshé, D. Nordli, P. Plouin, I. E. Scheffer, Revised terminology and concepts for organization of seizures and epilepsies: Report of the ILAE Commission on Classification and Terminology, 2005-2009. Epilepsia. 51, 676–685 (2010).

45. C. Helmstaedter, J. A. Witt, Epilepsy and cognition – A bidirectional relationship? Seizure. 49, 83–89 (2017).

46. C. Garcia-Ramos, J. J. Lin, T. S. Kellermann, L. Bonilha, V. Prabhakaran, B. P. Hermann, Graph theory and cognition: A complementary avenue for examining neuropsychological status in epilepsy. Epilepsy Behav. 64, 329–335 (2016).

47. M. Fukushima, R. F. Betzel, Y. He, M. A. de Reus, M. P. van den Heuvel, X. N. Zuo, O. Sporns, Fluctuations between high- and low-modularity topology in time-resolved functional connectivity. Neuroimage. 180, 406–416 (2018).

48. M. E. J. Newman, Modularity and community structure in networks. Proc. Natl. Acad. Sci. U. S. A. 103, 8577–8582 (2006).

49. M. Chavez, M. Valencia, V. Navarro, V. Latora, J. Martinerie, Functional modularity of background activities in normal and epileptic brain networks. Phys. Rev. Lett. 104 (2010), doi:10.1103/PhysRevLett.104.118701.

50. A. Goulas, R. F. Betzel, C. C. Hilgetag, Spatiotemporal ontogeny of brain wiring. Sci. Adv. 5 (2019), doi:10.1126/sciadv.aav9694.

51. G. Chen, Z. Duan, Network synchronizability analysis: A graph-theoretic approach. Chaos. 18, 1–10 (2008).

52. M. Barahona, L. M. Pecora, Synchronization in small-world systems. Phys. Rev. Lett. 89 (2002).

53. N. Meghanathan, Spectral Radius as a Measure of Variation in Node Degree for Complex Network Graphs. Proc. - 7th Int. Conf. u- e-Service, Sci. Technol. UNESST 2014, 30–33 (2015).

54. H. Chen, X. Zhao, F. Liu, S. Xu, W. Lu, Optimizing interconnections to maximize the spectral radius of interdependent networks. Phys. Rev. E. 95, 1–15 (2017).

55. Y. Wang, A. J. Trevelyan, A. Valentin, G. Alarcon, P. N. Taylor, M. Kaiser, Mechanisms underlying different onset patterns of focal seizures. PLoS Comput. Biol. 13, 1–22 (2017).

56. M. A. Kramer, E. D. Kolaczyk, H. E. Kirsch, Emergent network topology at seizure onset in humans. Epilepsy Res. 79, 173–186 (2008).

57. H. Blumenfeld, Cellular and network mechanisms of electrographic seizures. Epilepsia. 46, 21–33 (2005).

58. M. Rubinov, S. A. Knock, C. J. Stam, S. Micheloyannis, A. W. F. Harris, L. M. Williams, M. Breakspear, Small-world properties of nonlinear brain activity in schizophrenia. Hum. Brain Mapp. 30, 403–416 (2009).

59. C. H. Xia, Z. Ma, R. Ciric, S. Gu, R. F. Betzel, A. N. Kaczkurkin, M. E. Calkins, P. A. Cook, A. García de la Garza, S. N. Vandekar, Z. Cui, T. M. Moore, D. R. Roalf, K. Ruparel, D. H. Wolf, C. Davatzikos, R. C. Gur, R. E. Gur, R. T. Shinohara, D. S. Bassett, T. D. Satterthwaite, Linked dimensions of psychopathology and connectivity in functional brain networks. Nat. Commun. 9, 1–14 (2018).

60. T. D. Satterthwaite, E. Feczko, A. N. Kaczkurkin, D. A. Fair, Parsing Psychiatric Heterogeneity Through Common and Unique Circuit-Level Deficits. Biol. Psychiatry. 88, 4–5 (2020).

61. E. J. Cornblath, H. L. Li, L. Changolkar, B. Zhang, H. J. Brown, R. J. Gathagan, M. F. Olufemi, J. Q. Trojanowski, D. S. Bassett, V. M. Y. Lee, M. X. Henderson, Computational modeling of tau pathology spread reveals patterns of regional vulnerability and the impact of a genetic risk factor, 1–16 (2021).

62. W. G. Frankle, R. Narendran, Distinguishing Schizophrenia Subtypes: Can Dopamine Imaging Improve the Signal-to-Noise Ratio? Biol. Psychiatry. 87, 197–199 (2020).

63. S. L. Karalunas, J. T. Nigg, Heterogeneity and Subtyping in Attention-Deficit/Hyperactivity Disorder—Considerations for Emerging Research Using Person-Centered Computational Approaches. Biol. Psychiatry. 88, 103–110 (2020).

64. J. Dauwels, E. Eskandar, S. Cash, Localization of seizure onset area from intracranial non-seizure EEG by exploiting locally enhanced synchrony. Proc. 31st Annu. Int. Conf. IEEE Eng. Med. Biol. Soc. Eng. Futur. Biomed. EMBC 2009, 2180–2183 (2009).

65. A. R. Antony, A. V. Alexopoulos, J. A. González-Martínez, J. C. Mosher, L. Jehi, R. C. Burgess, N. K. So, R. F. Galán, Functional connectivity estimated from intracranial EEG predicts surgical outcome in intractable temporal lobe epilepsy. PLoS One. 8, 1–7 (2013).

66. G. Petkov, M. Goodfellow, M. P. Richardson, J. R. Terry, A critical role for network structure in seizure onset: A computational modeling approach. Front. Neurol. 5, 1–7 (2014).

67. D. J. Watts, S. H. Strogatz, Collective dynamics of “small-world” networks. Nature. 393, 440–442 (1998).

68. D. S. Bassett, E. Bullmore, Small-world brain networks. Neuroscientist. 12, 512–523 (2006).

69. F. V. Farahani, W. Karwowski, N. R. Lighthall, Application of graph theory for identifying connectivity patterns in human brain networks: A systematic review. Front. Neurosci. 13, 1–27 (2019).

70. J. P. Onnela, J. Saramäki, J. Kertész, K. Kaski, Intensity and coherence of motifs in weighted complex networks. Phys. Rev. E - Stat. Nonlinear, Soft Matter Phys. 71 (2005), doi:10.1103/PhysRevE.71.065103.

71. M. J. Vaessen, H. M. H. Braakman, J. S. Heerink, J. F. A. Jansen, M. H. J. A. Debeij-Van Hall, P. A. M. Hofman, A. P. Aldenkamp, W. H. Backes, Abnormal modular organization of functional networks in cognitively impaired children with frontal lobe epilepsy. Cereb. Cortex. 23, 1997–2006 (2013).

72. J. Reichardt, S. Bornholdt, Statistical mechanics of community detection. Phys. Rev. E - Stat. Nonlinear, Soft Matter Phys. 74, 1–14 (2006).

73. J. G. Restrepo, E. Ott, B. R. Hunt, Onset of synchronization in large networks of coupled oscillators. Phys. Rev. E - Stat. Nonlinear, Soft Matter Phys. 71, 1–12 (2005).

74. A. Jamakovic, R. E. Kooij, P. Van Mieghem, E. R. Van Dam, Robustness of networks against viruses: The role of the spectral radius. Proc. - 2006 Symp. Commun. Veh. Technol. IEEE SCVT 2006; 13th Annu. Symp. Commun. Veh. Technol. Benelux, 35–38 (2006).

75. Y. Wang, D. Chakrabarti, C. Wang, C. Faloutsos, Epidemic spreading in real networks: an eigenvalue viewpoint. 22nd Symp. Reliab. Distrib. Comput. Florence, Italy (2003).

76. E. R. van Dam, R. E. Kooij, The minimal spectral radius of graphs with a given diameter. Linear Algebra Appl. 423, 408–419 (2007).

77. R. Wang, Z. Z. Zhang, J. Ma, Y. Yang, P. Lin, Y. Wu, Spectral properties of the temporal evolution of brain network structure. Chaos. 25 (2015), doi:10.1063/1.4937451.

