## Supplementary information for "Brain network dynamics codify heterogeneity in seizure propagation"

### Supplementary Figures

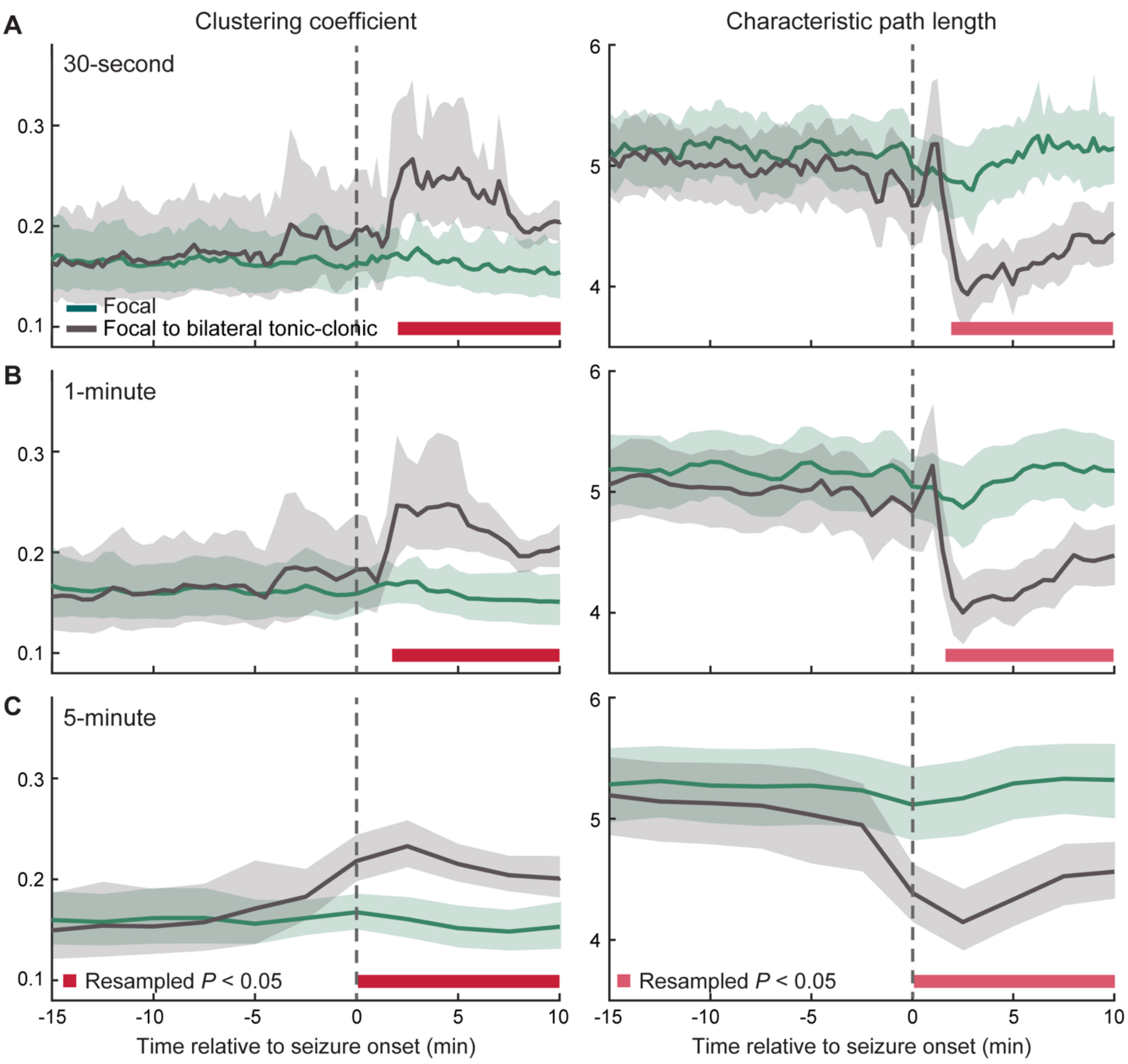

**Fig. S1. The effects of temporal smoothing on sample network measures.** Prior to computing graph theoretical features, a temporal smoothing parameter is first applied to the functional connectivity networks to approximate stationary pairwise connection strength signals. (**A**) The clustering coefficient and characteristic path length as computed after a smoothing parameter of 30 seconds has been applied to the adjacency matrices in a time-resolved fashion. These network measures are calculated separately for focal seizures that remain focal ($n$ = 49) and focal to bilateral tonic-clonic seizures ($n$ = 18). (**B**) Same network measures computed based on a temporal smoothing parameter of 1 minute. (**C**) Same network measures computed based on a temporal smoothing parameter of 5 minutes. The clustering coefficient of focal to bilateral tonic-clonic seizures are higher than that of focal seizures that remain focal (2-10 minutes; 2-10 minutes; seizure onset to 10 minutes after seizure onset for a smoothing parameter of 30 seconds, 1 minute, and 5 minutes, respectively). The characteristic path length of focal to bilateral tonic-clonic seizures are lower than that of focal seizures that remain focal (2-10 minutes; 2-10 minutes; seizure onset to 10 minutes after seizure onset for a smoothing parameter of 30 seconds, 1 minute, and 5 minutes, respectively). Error bars indicate 95% CIs computed by resampling the data distribution. Solid bars show resampled $P < 0.05$. All reported results in the main text are achieved based on a smoothing parameter of 30 seconds and the comparisons illustrated here suggest that our reported findings are robust and largely unaffected by the choice of this parameter.

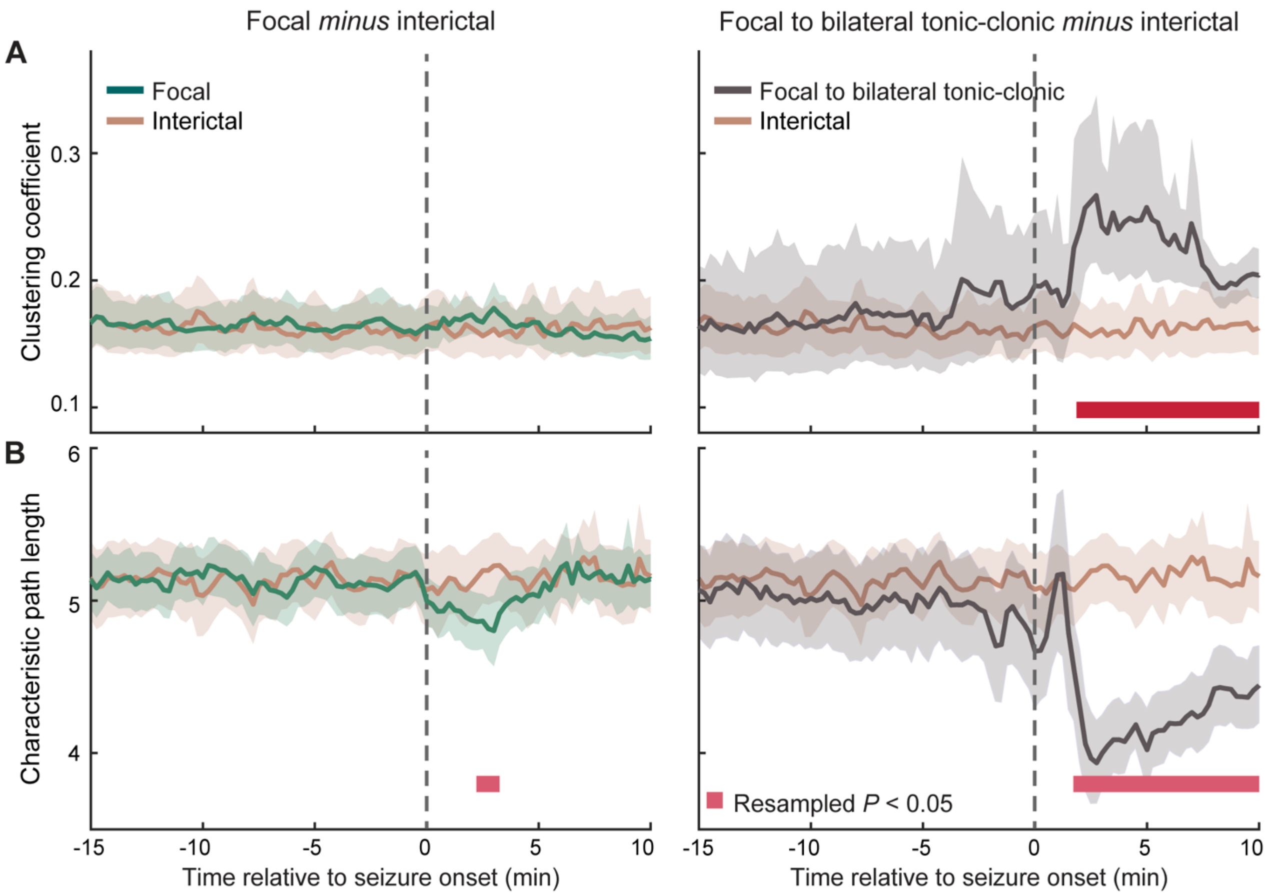


**Fig. S2. Small-world properties of focal seizures with constrained and unconstrained propagation dynamics relative to those of interictal periods.** (**A**) The clustering coefficient of focal seizures that remain focal ($n$ = 49) is not different from that of interictal networks ($n$ = 49). The clustering coefficient of focal to bilateral tonic-clonic seizures ($n$ = 18) is higher than that of interictal networks, 2-10 minutes after seizure onset. (**B**) The characteristic path length of focal seizures that remain focal is lower than that of interictal networks, 2.50-3.50 minutes after seizure onset. The characteristic path length of focal to bilateral tonic-clonic seizures is lower than that of interictal networks, 1.75-10 minutes after seizure onset. Error bars indicate 95% CIs computed by resampling the data distribution. Solid bars show resampled $P < 0.05$.

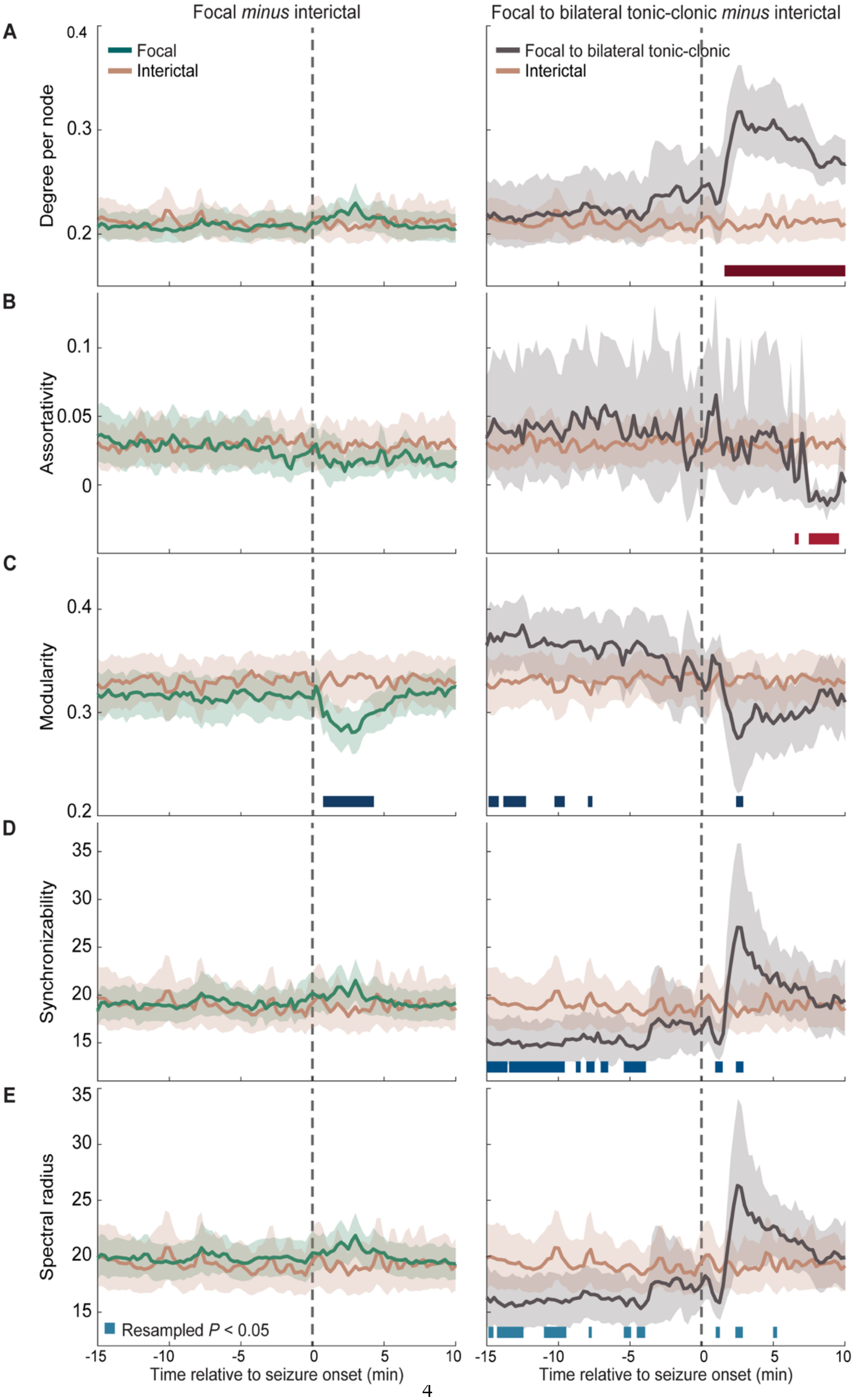

Focal minus interictal
Focal to bilateral tonic-clonic minus interictal
A
B
C
D
E
Focal
Interictal
Focal to bilateral tonic-clonic
Interictal
Degree per node
Assortativity
Modularity
Synchronizability
Spectral radius
Resampled P < 0.05
Time relative to seizure onset (min)
Time relative to seizure onset (min)

**Fig. S3. Network features of focal seizure with constrained and unconstrained dynamics relative to that of interictal periods. (A)** The degree per node of focal seizures that remain focal (*n* = 49) is not different from that of interictal networks (*n* = 49). The degree per of focal to bilateral tonic-clonic seizures (*n* = 18) is higher than that of interictal networks, 1.75-10 minutes after seizure onset. **(B)** The assortativity of focal seizures that remain focal is not different from that of inter-ictal networks. The assortativity of focal to bilateral tonic-clonic seizures is higher than that of interictal networks, 6.50-7 and 7.50-9.75 minutes after seizure onset. **(C)** The modularity of focal seizures that remain focal is lower than that of inter-ictal networks, 0.75-4.50 minutes after seizure onset. The modularity of focal to bilateral tonic-clonic seizures is higher than that of interictal networks, 14.75-14, 13.75-12, 10.25-9.50, and 7.75-7.50 minutes before seizure onset; and 2.50-3 minutes after onset. **(D)** The synchronizability of focal seizures that remain focal is not different from that of inter-ictal networks. The synchronizability focal to bilateral tonic-clonic seizures is lower than that of interictal networks, 15-13.5, 13.25-9.5, 8.75-8.25, 8-7.25, 7-6.50, and 5.25-3.75 minutes before seizure onset; and 1-1.5, and 2.50-3 minutes after seizure onset. **(E)** The spectral radius of focal seizures that remain focal is not different from that of interictal networks. The spectral radius of focal to bilateral tonic-clonic seizures is lower than that of interictal networks, 14.75-14.50, 14-12.25, 11-9.25, 7.75-7.50, 5.25-4.75, and 4.50-3.75 minutes before seizure onset; and 1-1.50, 2.50-3, and 5.25-5.50 minutes after seizure onset. Error bars indicate 95% CIs computed by resampling the data distribution. Solid bars show resampled $P < 0.05$.

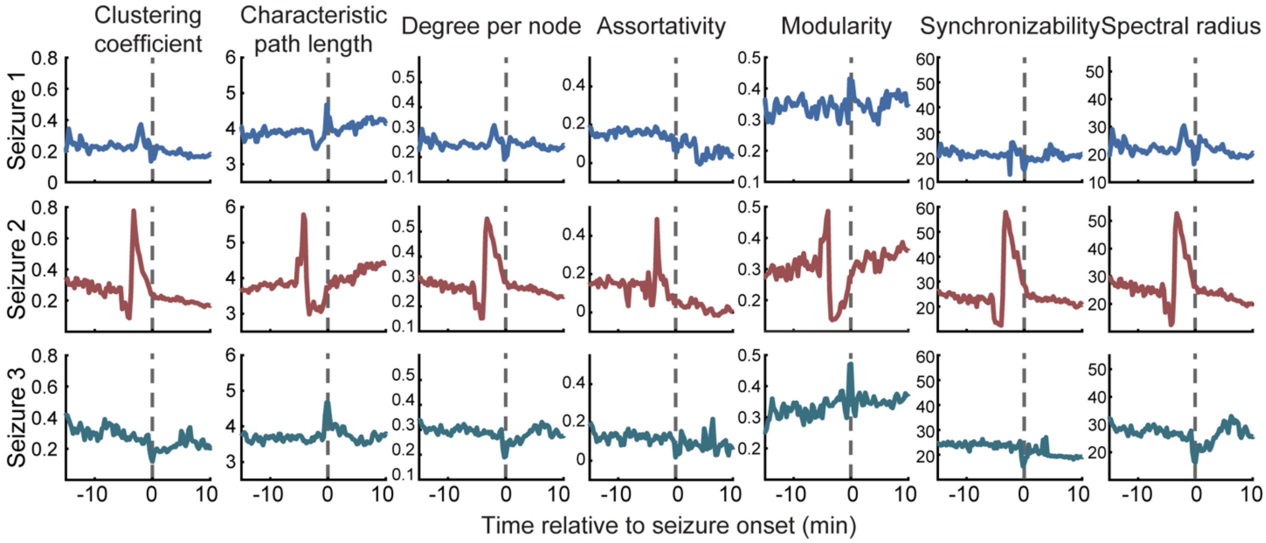


**Fig. S4. Distinct patterns of network properties across seizure types at a single-seizure level.** Graph theoretical measures extracted from networks associated with three seizures that share similar onset regions recorded from a sample patient. Seizure 1 and seizure 3 are categorized by an epileptologist as focal seizures that remain focal (sample recordings of seizure 1 is illustrated in Fig. 1B (left)), whereas seizure 2 is categorized as a focal to bilateral tonic-clonic seizure (sample recordings of seizure 2 is also illustrated in Fig. 1B (right)). Seizure 1 (top panel) and seizure 3 (bottom panel) exhibit similar patterns of topological properties, which differ from the features corresponding to seizure 2 (middle panel).
